# *In vitro* evaluation of the activity of pairwise combinations of zoliflodacin, gepotidacin, and ciprofloxacin against *Neisseria gonorrhoeae*

**DOI:** 10.64898/2026.07.07.737061

**Authors:** Bailey Bowcutt, Aditi Mukherjee, Samantha G. Palace, Yonatan H. Grad

## Abstract

Two new antibiotics, zoliflodacin and gepotidacin, were recently approved for the treatment of urogenital gonorrhea. While combination therapy could, in principle, delay the emergence and spread of resistance, doing so depends on the absence of antagonism between the co-administered drugs. Using *in vitro* checkerboard testing, we observed no evidence of antagonism for all pairwise combinations of zoliflodacin, gepotidacin, and ciprofloxacin, including in strains with elevated ciprofloxacin MICs.

## Introduction

*Neisseria gonorrhoeae* (*Ng*), the cause of the sexually transmitted infection gonorrhea, is a major public health problem due to a high burden of disease and increasing antibiotic resistance.^1,2^ Only one drug, ceftriaxone, remains recommended for empiric treatment,^2^ and ceftriaxone-resistant *N. gonorrhoeae* have been increasing in prevalence in Asia and spreading globally.^3–9^ Recently, two first-in-class drugs, zoliflodacin and gepotidacin, were approved by the United States Food and Drug Administration for treating uncomplicated urogenital gonorrhea.^10^ The optimal strategy for introducing the new antibiotics is not clear and depends on multiple factors.^11^ Combination therapy can slow the emergence of resistance under many scenarios^11^ and represents a strategy employed for other infectious diseases;^12–14^ moreover, for infections like gonorrhea, individuals may receive successive treatments such that the second antibiotic is taken before the first is fully cleared. As such, it is important to understand drug compatibility. Critically, if the two drugs are antagonistic, combination therapy, whether through co-administration or successive treatment, will be counterproductive.

Zoliflodacin and gepotidacin both target proteins within the *Ng* replisome, disrupting DNA topoisomerase activity and leading to cell death.^15,16^ Ciprofloxacin, which was used for gonorrhea treatment for many years, also targets the replisome,^17^ but has not been recommended for empiric gonorrhea treatment in the US since 2007 due to the high prevalence of resistance, conferred by target site mutations in *gyrA* and *parC*.^18^ However, *Ng* exposure to ciprofloxacin must still be considered, both because of the possibility of inappropriate treatment and because of bystander selection resulting from ciprofloxacin use for other indications. Gepotidacin and zoliflodacin are active against ciprofloxacin-resistant *Ng*.^19,20^ Nevertheless, the target sites of these drugs are in close spatial proximity to one another,^21,22^ and their shared mechanism of action raises the possibility that administering these drugs in combination could either enhance or reduce overall antibacterial efficacy. Determining how these replisome-targeting drugs interact with each other to inhibit *Ng* growth, and how these interactions are modulated by existing ciprofloxacin-resistance mutations, could help inform clinical considerations around combination therapy.

A previous study evaluated interactions between zoliflodacin and ciprofloxacin in a panel of *Ng* isolates, reporting synergy between these drugs for two mutant strains harboring the zoliflodacin resistance mutation *gyrB*^D429N^, but reporting largely additive interactions in clinical isolates with no evidence of antagonism.^23^ It is not yet known how consistent these results will be in the presence of important ciprofloxacin resistance-associated *gyrA* mutations, or across strains from more diverse genetic backgrounds representative of epidemiologically significant gonococcal lineages. Gepotidacin has been tested for synergy and antagonism with a range of drugs, but not in combination with ciprofloxacin or zoliflodacin in *Ng*. Interestingly, synergy was observed between gepotidacin and moxifloxacin for one out of three *Ng* strains tested in a checkerboard analysis.^24^ However, the limited number of strains in this study did not include common resistance-associated mutations that have arisen in response to ciprofloxacin use, and no studies have examined the interaction between gepotidacin and zoliflodacin.

Here, we test pairwise combinations of zoliflodacin, gepotidacin, and ciprofloxacin against a panel of 16 genetically diverse *Ng* strains that include representatives of clinically important ciprofloxacin resistance-associated alleles in *gyrA* and *parC*. To test the hypothesis that target site resistance-associated mutations might modulate these interactions, we also tested the same drug combinations for synergy or antagonism in an isogenic panel of strains harboring ciprofloxacin resistance-associated mutations in *gyrA* at amino acid positions 91 and 95 and the *gyrB* substitution D429N, which can increase resistance to both ciprofloxacin and zoliflodacin.^25^

## Materials and Methods

### Phylogenetic Tree and Isolate Selection

A panel of 15 strains was selected to represent a variety of strain backgrounds across a phylogenetic tree of 742 isolates within our laboratory’s collection. The laboratory strain MS11 was added to this panel as a representative of the wildtype *gyrA*^91S/95D^ ciprofloxacin-sensitive genetic background. Pseudogenome alignments with recombinant regions masked were generated with Gubbins^26^, and a phylogenetic tree was built using IQTREE v2.4.0^27^ using ModelFinder^28^ to select the optimal substitution model. For the strain H18-208, only long-read sequencing data were available, so short reads were simulated by ART v2016.06.05.^29^ The phylogeny was visualized using iTOL v7.^30^

### Bacterial Isolates and Culture Conditions

*N. gonorrhoeae* strains were cultured on GCB agar (Difco) with Kellogg’s supplements (GCB-K)^31^ at 37°C in 5% CO_2_ atmospheric conditions.

Strains are listed in Supplementary File 1. A panel of isogenic strains with resistance-associated *gyrA* and *gyrB* mutations was created in the clinical isolate GCGS0481^32^, which natively harbors the *gyrA* ^91F/95G^ allele. To introduce alternative *gyrA* alleles into this strain, the ciprofloxacin-susceptible allele GyrA^91S/95D^ was introduced with a kanamycin marker as described.^25^ Alleles containing resistance mutations in *gyrA* positions 91 and 95 were then amplified as described^33^ and introduced into GCGS0481 GyrA^91S/95D^ via electroporation and selection on GCB-K with 2 µg/mL ciprofloxacin (for alleles containing *gyrA*^S91F^) or 0.25 µg/mL ciprofloxacin (for alleles with substitutions at *gyrA*^95^ only). There was no kanamycin marker in the GyrA^91F/95A^ strain due to recombination occurring before the marker. Expected MICs did not vary in strains with/without kanamycin markers. Similarly, *gyrB*^D429N^ was amplified as described^33^ and introduced via electroporation and selection on GCB-K with a dried droplet of 4 µg/mL zoliflodacin. Transformant genotypes were verified by Sanger sequencing as described.^33^

### Agar dilution and checkerboard assay

MICs were measured via agar dilution. For drug interaction testing, drugs were combined pairwise in a six-by-six checkerboard format with agar poured into 24-well culture plates. Doubling dilutions of each drug were tested, with concentrations ranging from 0.125x MIC to 2x MIC for each strain. The pairwise drug combinations tested were zoliflodacin with gepotidacin, zoliflodacin with ciprofloxacin, and gepotidacin with ciprofloxacin.

Cultures were grown for MIC testing for 18-20 hours on GCB-K agar as above, after which a barely turbid suspension (OD ∼0.1) was prepared in tryptic soy broth. A 2 μL spot of this suspension was plated on agar in each well of the checkerboard plates and read after 24 hours of growth at 37°C in 5% CO_2_. Two biological replicates were performed for each test.

### Interpretation

The fractional inhibitory concentration index (FICI) was calculated as described^34^ to determine if synergy or antagonism occurred, with the following breakpoints: synergy, FICI ≤0.5; indifference, FICI > 0.5–4.0; and antagonism, FICI > 4.0.

## Results

To examine interactions between replisome-targeting drugs in phylogenetically diverse *Ng*, we selected a panel of isolates to represent a variety of genetic backgrounds, comprising 15 recent clinical isolates and the laboratory strain MS11 (Supplementary File 2). All 15 clinical isolates carried the ciprofloxacin resistance variant *gyrA*^S91F^, reflecting the high prevalence of this allele in clinical isolates. Isolates were selected to contain substitutions at *gyrA*^95^ (D95G/A/N/Y), reflecting the allelic variation at this position among ciprofloxacin-resistant *N. gonorrhoeae*.

Furthermore, because *parC* substitutions are known to contribute to resistance to both ciprofloxacin and gepotidacin, we selected isolates containing variants at the polymorphic positions 86, 87, and 91.^15^ Genotypic information for all isolates is available in Supplementary Table 3.

Drug interactions between pairwise combinations of zoliflodacin, gepotidacin, and ciprofloxacin for this panel are summarized in Table 1. All drug interactions were in the indifferent category for all isolates. We observed no antagonism.

**Table 1.** Panel checkerboard drug interactions. The mean fractional inhibitory concentration index (FICI) of pairwise drug combinations of zoliflodacin (ZFD), gepotidacin (GEP), and ciprofloxacin (CIP) of 15 clinical samples and one ciprofloxacin-sensitive laboratory sample. The mean was calculated including all FICIs across the plates, while the range represents the FICIs that appeared in individual wells. Two biological replicates were performed for each test.

| Isolate name | Mean FICI (Range) |  |  |
| --- | --- | --- | --- |
|  | ZFD X GEP | GEP X CIP | ZFD X CIP |
| CCC033 | 0.793(0.567-1.128) | 0.906(0.625-1.250) | 0.916(0.633-1.128) |
| DDD033 | 1.055(0.633-1.254) | 0.896(0.625-1.125) | 0.968(0.754-1.256) |
| EEE016 | 0.865(0.628-1.128) | 0.945(0.624-1.250) | 1.054(0.754-1.250) |
| EEE036 | 0.866(0.632-1.127) | 1.042(0.624-1.252) | 1.267(1.008-1.500) |
| GCGS0860 | 0.874(0.624-1.124) | 1.087(0.750-1.250) | 1.054(0.756-1.250) |
| H18-208 | 0.790(0.498-1.250) | 1.136(1.000-1.252) | 1.355(1.008-2.109) |
| HHH012 | 1.099(0.754-1.250) | 0.906(0.625-1.250) | 0.914(0.754-1.128) |
| HHH014 | 0.692(0.563-1.063) | 1.091(0.626-1.500) | 0.959(0.627-1.250) |
| HHH023 | 0.663(0.506-1.064) | 1.047(0.750-1.250) | 1.054(0.756-1.250) |
| HHH040 | 1.102(0.754-1.254) | 1.200(0.752-2.125) | 0.965(0.754-1.128) |
| MS11 | 0.974(0.625-1.250) | 0.966(0.563-1.250) | 1.144(1.004-1.256) |
| Ng175 | 0.707(0.504-1.062) | 1.138(1.004-1.256) | 1.144(1.000-1.250) |
| Ng183 | 1.100(0.754-1.256) | 1.056(0.750-1.250) | 1.153(0.754-1.508) |
| NY0215 | 0.977(0.566-1.254) | 0.813(0.626-1.125) | 0.727(0.506-1.127) |
| NY0738 | 0.977(0.567-1.254) | 1.008(0.562-1.500) | 1.039(0.754-1.254) |

Zoliflodacin, gepotidacin, and ciprofloxacin have overlapping drug targets, and some cross-resistance between these drugs has been observed in *Ng*.^21,25^ To test whether individual resistance mutations affect drug interactions between zoliflodacin, gepotidacin, and ciprofloxacin, we measured pairwise drug interactions in an isogenic strain panel in which common ciprofloxacin resistance-associated mutations were added at *gyrA* positions 91 and 95 alone or in combination with the zoliflodacin resistance mutation *gyrB*^D429N^ in a clinical isolate, GCGS0481 (Table 2). Almost all drug combinations were in the indifferent category for all isogenic strains; while a single replicate, the GCGS0481 *gyrA*^91S/95N^ *gyrB*^D429N^ strain showed synergy (FICI 0.37) with inhibition at a zoliflodacin concentration of 2 mg/L in combination with a gepotidacin concentration of 0.25 mg/L, this result was not consistent across experimental replicates. We observed no antagonism.

**Table 2.** GCGS0481 isogenic strain panel checkerboard drug interactions. The fractional inhibitory concentration index (FICI) of pairwise drug combinations of zoliflodacin (ZFD), gepotidacin (GEP), and ciprofloxacin (CIP) in an isogenic strain panel using isolate GCGS0481 and relevant resistance-associated mutations. The mean was calculated including all FICIs across the plates, while the range represents the FICIs that appeared in individual wells. Two biological replicates were performed for each test.

| Strain name | Mean FICI (range) |  |  |
| --- | --- | --- | --- |
|  | ZFD X GEP | GEP X CIP | ZFD X CIP |
| GyrA<sup>91F/95G</sup> | 1.050 (0.748-1.248) | 0.969 (0.624-1.250) | 1.189 (0.750-1.504) |
| GyrA<sup>91F/95G</sup> GyrB<sup>D429N</sup> | 1.006 (0.624-1.250) | 1.036 (0.750-1.252) | 1.094 (0.750-1.250) |
| GyrA<sup>91F/95A</sup> | 1.056 (0.628-2.124) | 1.006 (0.750-1.250) | 1.099 (0.756-1.252) |
| GyrA<sup>91F/95A</sup> GyrB<sup>D429N</sup> | 0.993 (0.625-1.250) | 1.135 (1.000-1.250) | 0.854 (0.562-1.125) |
| GyrA<sup>91F/95D</sup> | 1.093 (0.752-1.250) | 1.097 (0.750-1.250) | 0.967 (0.562-1.252) |
| GyrA<sup>91F/95D</sup> GyrB<sup>D429N</sup> | 1.142 (1.000-1.250) | 0.986 (0.750-1.250) | 0.863 (0.625-1.125) |
| GyrA<sup>91F/95N</sup> | 1.094 (0.752-1.252) | 0.974 (0.626-1.250) | 1.239 (1.000-1.504) |
| GyrA<sup>91F/95N</sup> GyrB<sup>D429N</sup> | 0.999 (0.748-1.250) | 0.965 (0.750-1.250) | 1.011 (0.750-1.250) |
| GyrA<sup>91S/95G</sup> | 1.050 (0.748-1.250) | 0.977 (0.748-1.252) | 1.007 (0.628-1.250) |
| GyrA<sup>91S/95G</sup> GyrB<sup>D429N</sup> | 1.099 (0.748-1.250) | 0.840 (0.625-1.126) | 1.094 (0.750-1.250) |
| GyrA<sup>91S/95A</sup> | 1.006 (0.748-1.248) | 0.898 (0.502-1.254) | 1.305 (1.004-2.128) |
| GyrA<sup>91S/95A</sup> GyrB<sup>D429N</sup> | 0.862 (0.624-1.125) | 0.950 (0.562-1.252) | 1.602 (0.750-2.500) |
| GyrA<sup>91S/95D</sup> | 1.050 (0.748-1.248) | 1.097 (0.624-1.500) | 0.896 (0.625-1.126) |
| GyrA<sup>91S/95D</sup> GyrB<sup>D429N</sup> | 1.149 (1.000-1.250) | 1.153 (1.000-1.252) | 0.723 (0.504-1.127) |
| GyrA<sup>91S/95N</sup> | 0.965 (0.624-1.250) | 1.091 (0.750-1.252) | 0.972 (0.750-1.126) |
| GyrA<sup>91S/95N</sup> GyrB<sup>D429N</sup> | 0.609 (0.374-0.750) | 0.785 (0.500-1.126) | 1.297 (1.000-2.125) |

## Discussion

We investigated the effects of zoliflodacin, gepotidacin, and ciprofloxacin in pairwise combinations and found indifferent or additive drug interactions across a range of drug combinations. This pattern was consistent across 16 isolates with a range of resistance profiles and *gyrA, gyrB*, and *parC* alleles, as well as an isogenic panel with clinically relevant mutations in *gyrA* and *gyrB*. Resistance alleles did not alter interaction outcomes. Confirming the absence of antagonism is relevant given the possibility that treatment efficacy may be impacted if other antibiotics were recently taken and not yet cleared, and for consideration of the possibility of combination therapy to suppress the emergence of resistance across *N. gonorrhoeae* genetic backgrounds.^35–37^

There is substantial interest in understanding drug interactions and in developing possible combination therapies for *Ng*, both for directed treatment and for empiric and directed treatment of *Ng* coinfection with *Chlamydia trachomatis*. Nonsynergistic combinations have been used for *Ng* treatment; for example, the ceftriaxone and azithromycin combination was indifferent or additive in 90/95 strains tested.^35^ For possible coinfection with *C. trachomatis*, doxycycline is often given empirically,^2^ motivating interest in testing the efficacy of new gonorrhea therapeutics when in combination with doxycycline. The combination of doxycycline and zoliflodacin has had mixed results in various *in vitro* assays, with an indifferent effect in checkerboard assays and a negative effect of tetracycline with zoliflodacin on bacterial killing in a 6-hour time kill assay.^38^ This interaction emphasizes the importance of checking for antagonism in each drug pair. However, in the more biologically relevant hollow fiber infection model, the combination of zoliflodacin and doxycycline was efficacious and superior at suppressing resistance compared to zoliflodacin monotherapy against *Ng*.^39^ The combination of gepotidacin with doxycycline has not been explored for *Ng*, but preliminary data indicate that this combination may show synergy against *Mycoplasma genitalium*.^40^ Increasing rates of doxycycline resistance in *Ng* in the US and already high prevalence in many other locations may render moot considerations of doxycycline-based combinations for directed *Ng* treatment.^41–43^

In conclusion, zoliflodacin, gepotidacin, and ciprofloxacin have largely indifferent interactions in diverse clinical *Ng* isolates, and this indifference is maintained in the presence of key resistance mutations and in genetic backgrounds with high-level ciprofloxacin resistance. Further work to determine the viability of combination therapy includes *in vivo* drug interaction testing, safety testing, and studies to determine the effects of multiple drugs on the microbiome. Importantly, the lack of antagonism among these drug classes offers a measure of reassurance for consideration of their combination or overlapping use in clinical contexts.

## Author contributions

BB, SGP, and YHG conceptualized the study. AM and BB acquired/analyzed the data. BB, SGP, and YHG wrote the original draft, and all the authors reviewed and edited the manuscript.

## Acknowledgments

We would like to thank the Grad lab for their helpful feedback on this work. This work was supported by NIH R01 AI132606 and R01 AI153521 grants to YHG. This material is based upon work supported by the National Science Foundation Graduate Research Fellowship Program under Grant No. DGE 2140743 to BB. Any opinions, findings, conclusions, or recommendations expressed in this material are those of the author(s) and do not necessarily reflect the views of the National Science Foundation or the National Institutes of Health.

---

**Supplementary File 1.**
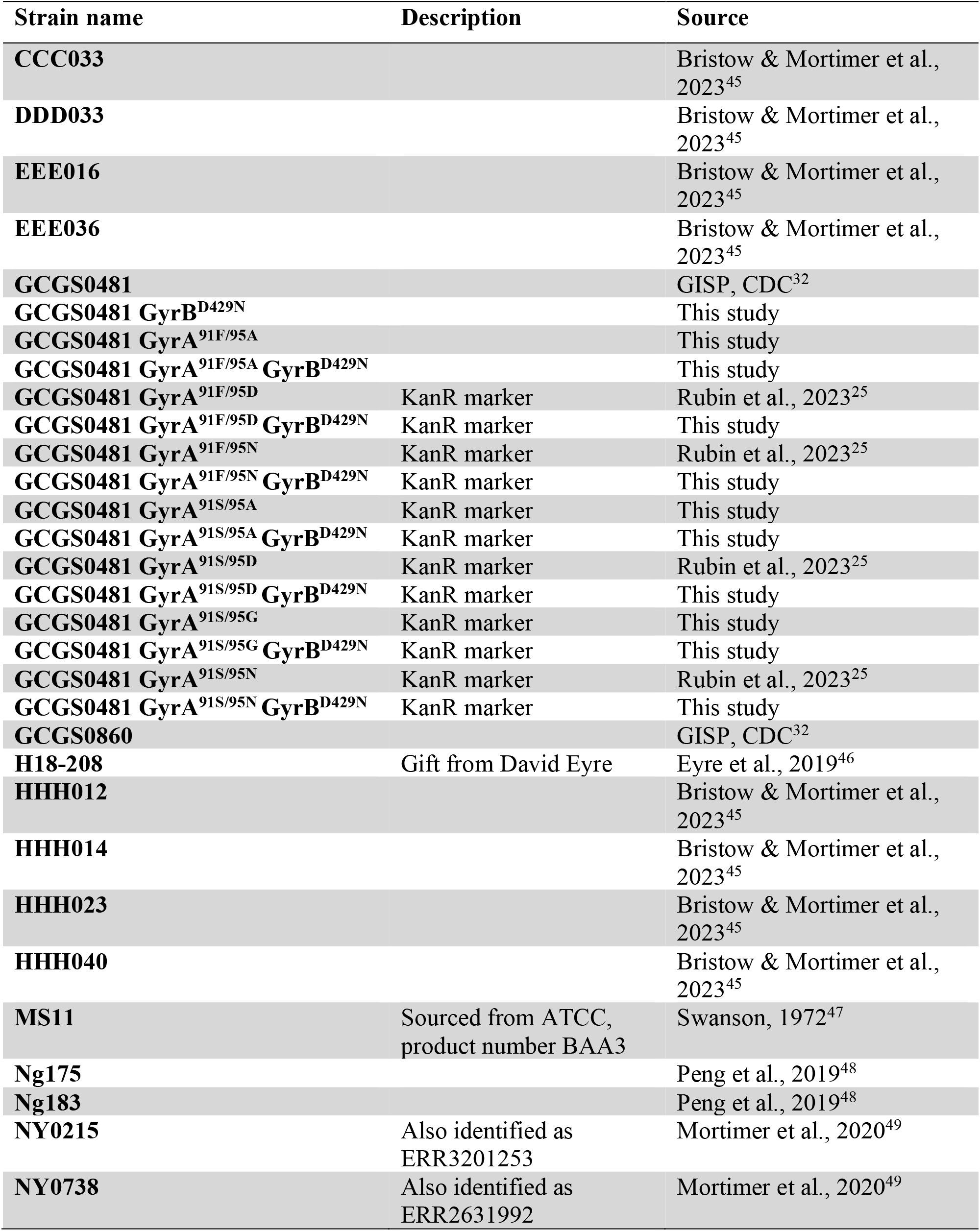
*N. gonorrhoeae* strains and primers used in this study.

**Supplementary file 2.**
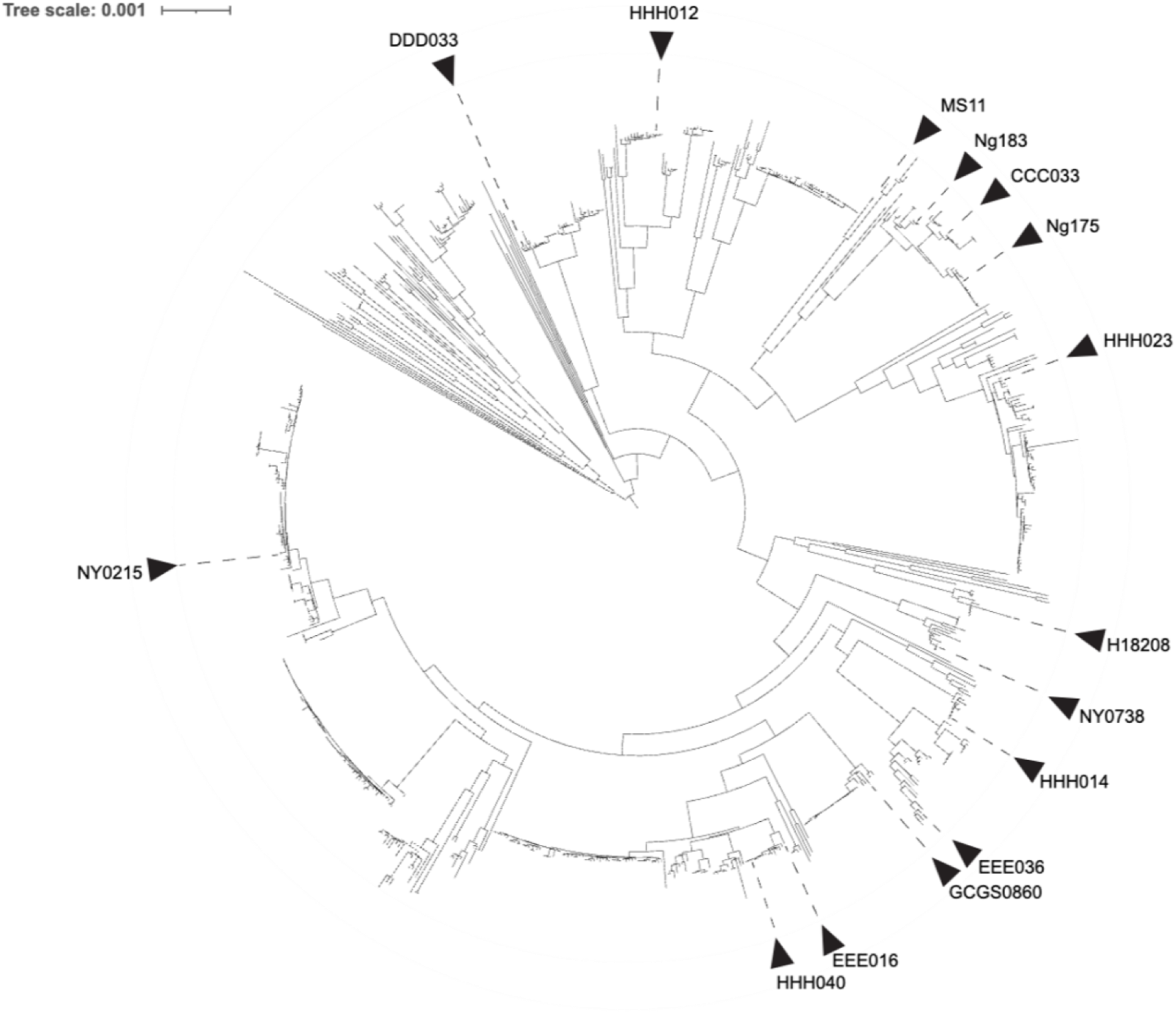
Phylogenetic tree. n = 742. Pseudogenome alignments with recombinant regions masked were generated with Gubbins^26^, and a phylogenetic tree was built using IQTREE v2.4.0^27^ using ModelFinder^28^ to select the optimal substitution model. For the strain H18-208, only long-read sequencing data were available, so short reads were simulated by ART v2016.06.05.^29^ The phylogeny was visualized using iTOL v7.^30^

**Supplementary file 3.**
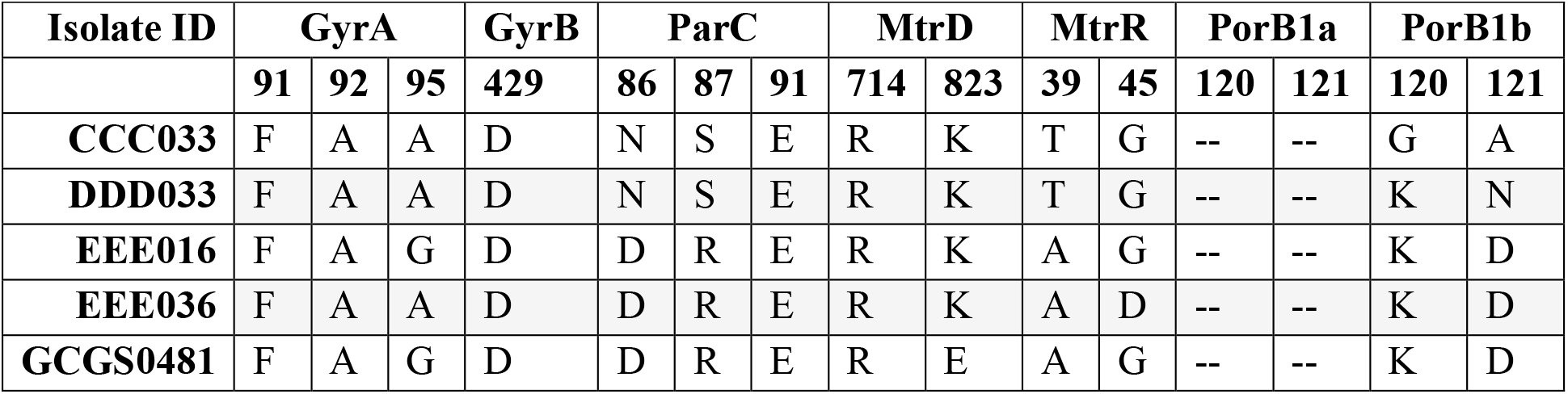
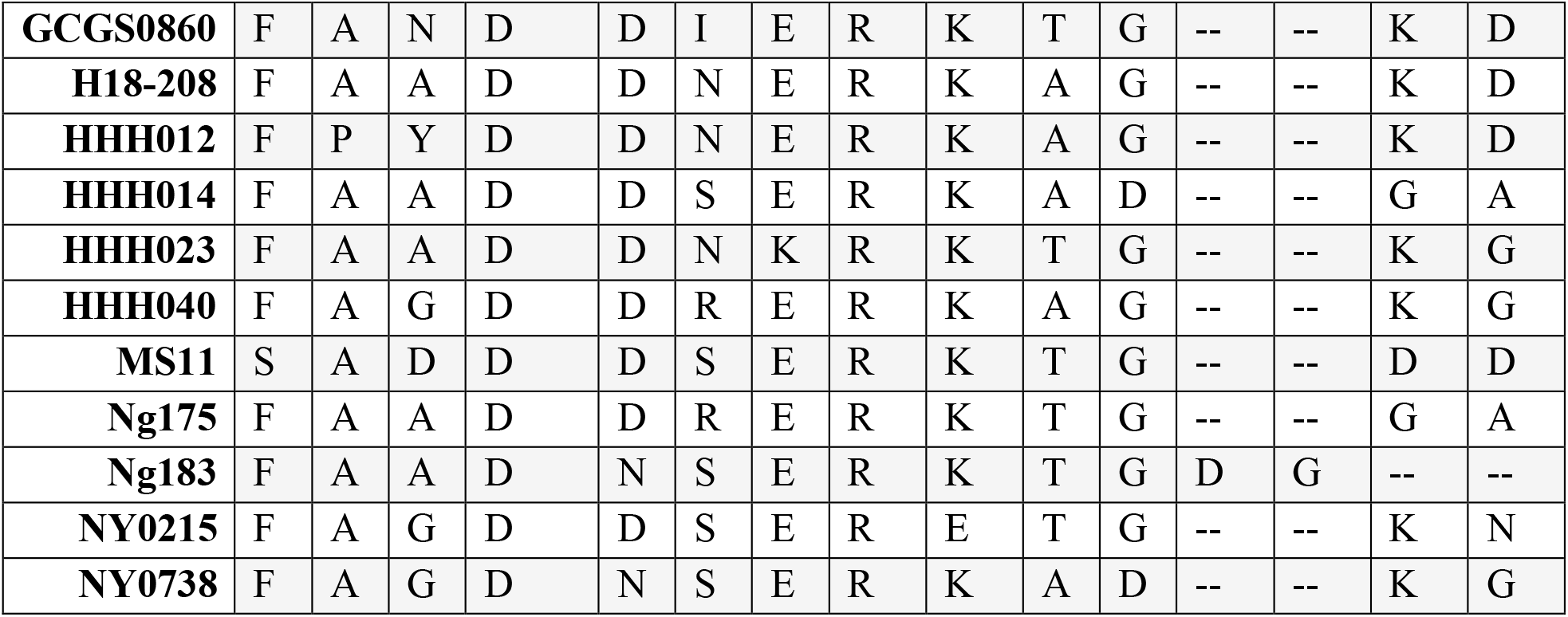
Strain metadata of major resistance alleles.

**Supplementary File 4.**
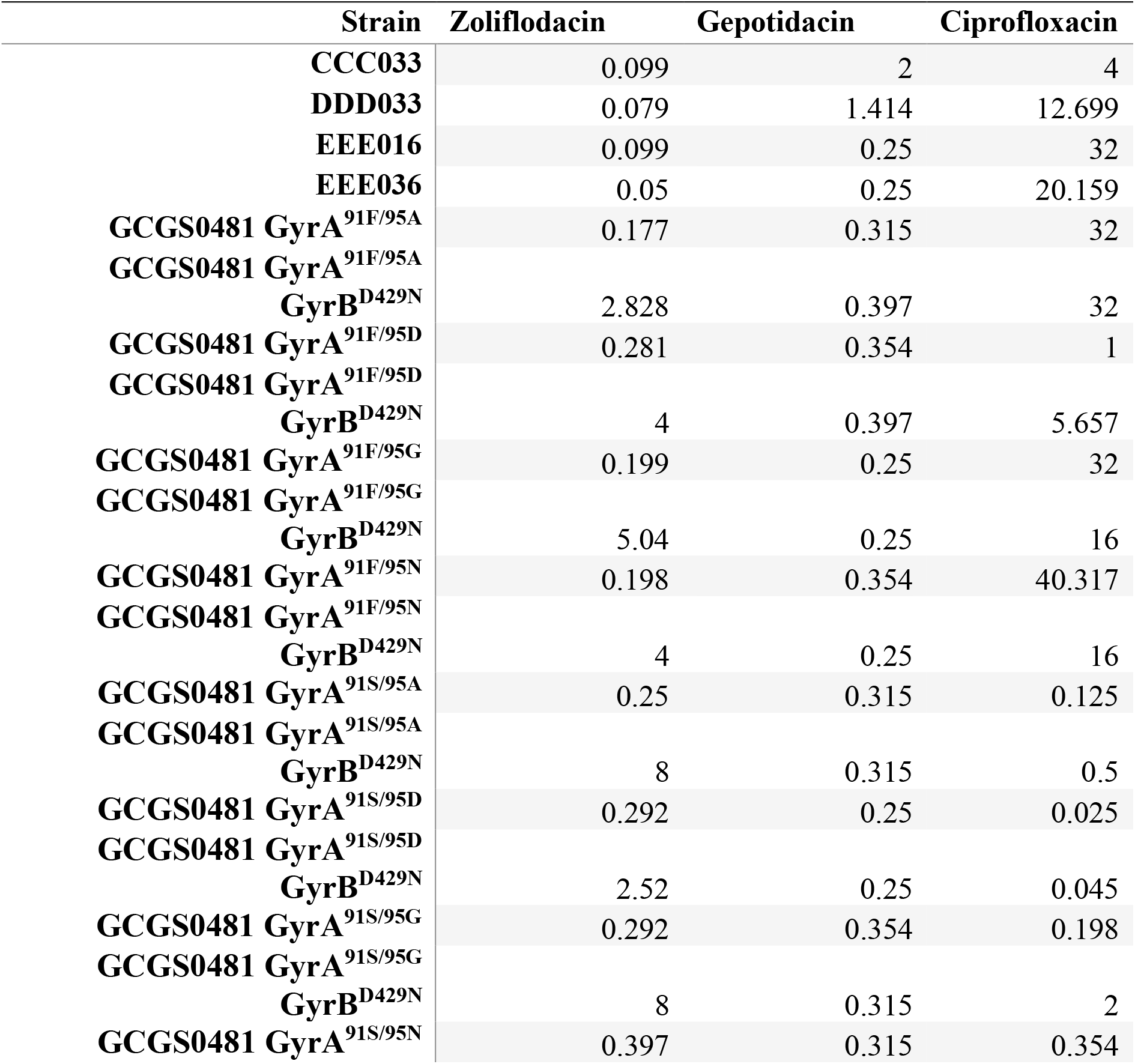
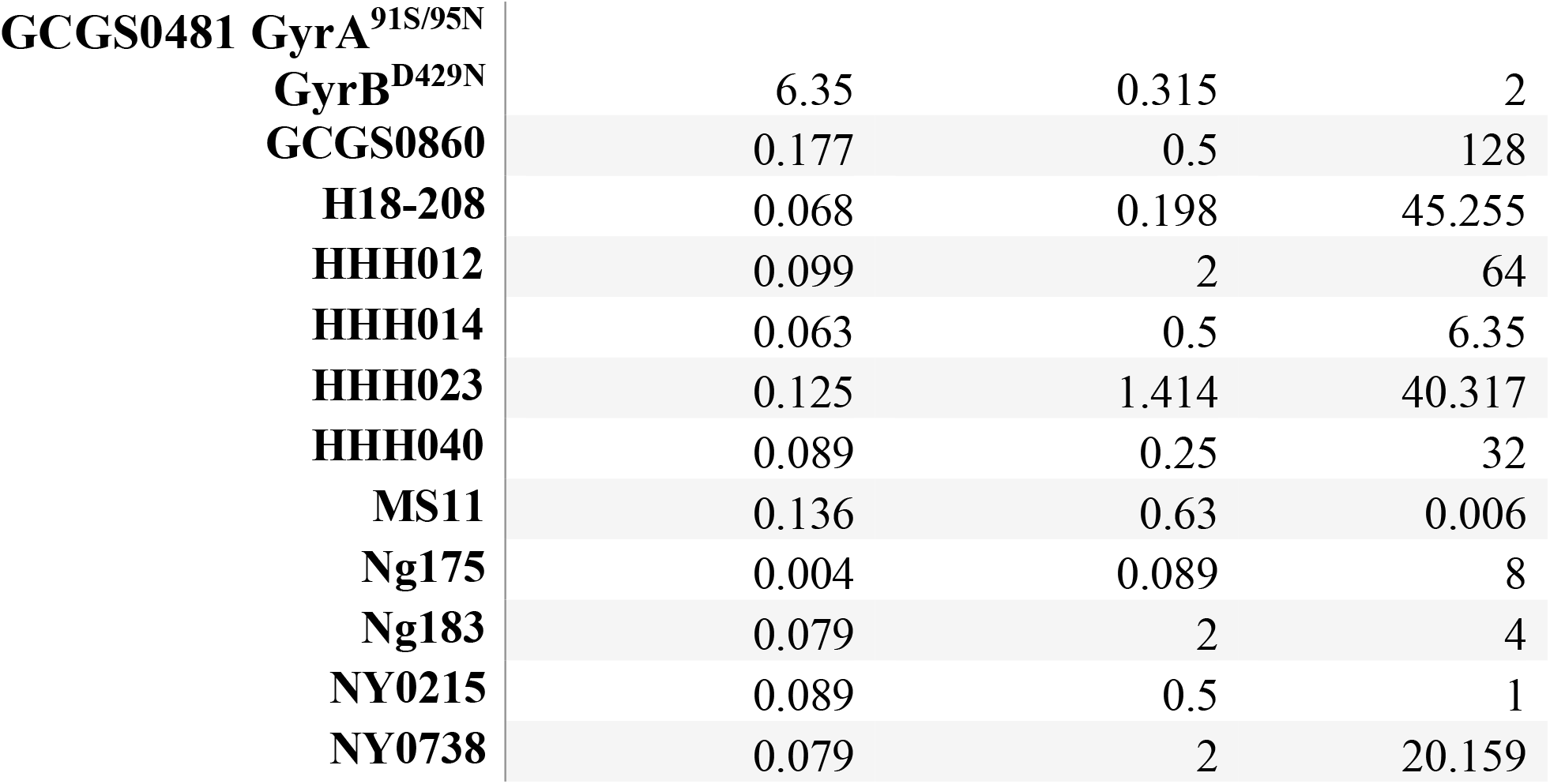
Geometric Mean MIC across all replicates (*n*=4 in each cell)

## References

1. Centers for Disease Control and Prevention (CDC). Sexually Transmitted Infections Surveillance, 2024 (Provisional). STI Statistics https://www.cdc.gov/sti-statistics/annual/index.html (2025).

2. Centers for Disease Control and Prevention (CDC). Sexually Transmitted Infections Treatment Guidelines, 2021. https://www.cdc.gov/std/treatment-guidelines/gonorrhea-adults.htm.

3. Berçot, B. et al. Ceftriaxone-resistant, multidrug-resistant Neisseria gonorrhoeae with a novel mosaic penA-237.001 gene, France, June 2022. Eurosurveillance 27, 2200899 (2022).

4. Fifer, H. et al. Ceftriaxone-resistant Neisseria gonorrhoeae detected in England, 2015–24: an observational analysis. J. Antimicrob. Chemother. 79, 3332–3339 (2024).

5. Day, M. et al. Detection of 10 cases of ceftriaxone-resistant Neisseria gonorrhoeae in the United Kingdom, December 2021 to June 2022. Eurosurveillance 27, 2200803 (2022).

6. Adamson, P. C., Hieu, V. N., Nhung, P. H., Whiley, D. M. & Chau, T. M. Ceftriaxone resistance in Neisseria gonorrhoeae associated with the penA-60.001 allele in Hanoi, Viet Nam. Lancet Infect. Dis. 24, e351–e352 (2024).

7. Laumen, J. G. E. et al. High Prevalence of Ceftriaxone-Resistant Neisseria gonorrhoeae in Hanoi, Vietnam, 2023–2024. J. Infect. Dis. 232, e73–e77 (2025).

8. Ouk, V. et al. High prevalence of ceftriaxone-resistant and XDR Neisseria gonorrhoeae in several cities of Cambodia, 2022–23: WHO Enhanced Gonococcal Antimicrobial Surveillance Programme (EGASP). JAC-Antimicrob. Resist. 6, dlae053 (2024).

9. Zhu, X., Xi, Y., Gong, X. & Chen, S. Ceftriaxone-Resistant Gonorrhea — China, 2022. Morb. Mortal. Wkly. Rep. 73, 255–259 (2024).

10. U.S. Food and Drug Administration. FDA Approves Two Oral Therapies to Treat Gonorrhea. U.S. Food and Drug Administration https://www.fda.gov/news-events/press-announcements/fda-approves-two-oral-therapies-treat-gonorrhea (2025).

11. Reichert, E. et al. Resistance-minimising strategies for introducing a novel antibiotic for gonorrhoea treatment: a mathematical modelling study. Lancet Microbe 4, e781–e789 (2023).

12. Tyers, M. & Wright, G. D. Drug combinations: a strategy to extend the life of antibiotics in the 21st century. Nat. Rev. Microbiol. 17, 141–155 (2019).

13. WHO consolidated guidelines on tuberculosis: module 4: treatment and care. https://www.who.int/publications/i/item/9789240107243.

14. WHO updated recommendations on HIV clinical management: recommendations for a public health approach. https://www.who.int/publications/i/item/9789240119468.

15. Oviatt, A. A. et al. Mechanism of Action of Gepotidacin: Well-Balanced Dual-Targeting against Neisseria gonorrhoeae Gyrase and Topoisomerase IV in Cells and In Vitro. ACS Infect. Dis. 11, 3093–3105 (2025).

16. Alm, R. A. et al. Characterization of the Novel DNA Gyrase Inhibitor AZD0914: Low Resistance Potential and Lack of Cross-Resistance in Neisseria gonorrhoeae. Antimicrob. Agents Chemother. 59, 1478–1486 (2015).

17. Collins, J. A., Oviatt, A. A., Chan, P. F. & Osheroff, N. Target-Mediated Fluoroquinolone Resistance in Neisseria gonorrhoeae: Actions of Ciprofloxacin against Gyrase and Topoisomerase IV. ACS Infect. Dis. 10, 1351–1360 (2024).

18. Centers for Disease Control and Prevention (CDC). Update to CDC’s sexually transmitted diseases treatment guidelines, 2006: fluoroquinolones no longer recommended for treatment of gonococcal infections. MMWR Morb. Mortal. Wkly. Rep. 56, 332–336 (2007).

19. Ross, J. D. C. et al. Oral gepotidacin for the treatment of uncomplicated urogenital gonorrhoea (EAGLE-1): a phase 3 randomised, open-label, non-inferiority, multicentre study. The Lancet 405, 1608–1620 (2025).

20. Bradford, P. A., Miller, A. A., O’Donnell, J. & Mueller, J. P. Zoliflodacin: An Oral Spiropyrimidinetrione Antibiotic for the Treatment of Neisseria gonorrheae, Including Multi-Drug-Resistant Isolates. ACS Infect. Dis. 6, 1332–1345 (2020).

21. Mukherjee, A. et al. Genetic Background Modulates Zoliflodacin and Gepotidacin Cross-Resistance and Fitness in Neisseria gonorrhoeae. J. Infect. Dis. jiag174 (2026) doi:10.1093/infdis/jiag174.

22. Scangarella-Oman, N. E. et al. Microbiological Analysis from a Phase 2 Randomized Study in Adults Evaluating Single Oral Doses of Gepotidacin in the Treatment of Uncomplicated Urogenital Gonorrhea Caused by Neisseria gonorrhoeae. Antimicrob. Agents Chemother. 62, 10.1128/aac.01221-18 (2018).

23. Foerster, S. et al. Genetic Resistance Determinants, In Vitro Time-Kill Curve Analysis and Pharmacodynamic Functions for the Novel Topoisomerase II Inhibitor ETX0914 (AZD0914) in Neisseria gonorrhoeae. Front. Microbiol. 6, (2015).

24. Farrell, D. J., Sader, H. S., Rhomberg, P. R., Scangarella-Oman, N. E. & Flamm, R. K. In Vitro Activity of Gepotidacin (GSK2140944) against Neisseria gonorrhoeae. Antimicrob. Agents Chemother. 61, 10.1128/aac.02047-16 (2017).

25. Rubin, D. H., Mortimer, T. D. & Grad, Y. H. Neisseria gonorrhoeae diagnostic escape from a gyrA-based test for ciprofloxacin susceptibility and the effect on zoliflodacin resistance: a bacterial genetics and experimental evolution study. Lancet Microbe 4, e247– e254 (2023).

26. Croucher, N. J. et al. Rapid phylogenetic analysis of large samples of recombinant bacterial whole genome sequences using Gubbins. Nucleic Acids Res. 43, e15 (2015).

27. Minh, B. Q. et al. IQ-TREE 2: New Models and Efficient Methods for Phylogenetic Inference in the Genomic Era. Mol. Biol. Evol. 37, 1530–1534 (2020).

28. Kalyaanamoorthy, S., Minh, B. Q., Wong, T. K. F., von Haeseler, A. & Jermiin, L. S. ModelFinder: fast model selection for accurate phylogenetic estimates. Nat. Methods 14, 587–589 (2017).

29. Huang, W., Li, L., Myers, J. R. & Marth, G. T. ART: a next-generation sequencing read simulator. Bioinformatics 28, 593–594 (2012).

30. Letunic, I. & Bork, P. Interactive Tree of Life (iTOL) v6: recent updates to the phylogenetic tree display and annotation tool. Nucleic Acids Res. 52, W78–W82 (2024).

31. Kellogg, D. S., Peacock, W. L., Deacon, W. E., Brown, L. & Pirkle, C. I. Neisseria gonorrhoeae I: Virulence Genetically Linked to Clonal Variation. J. Bacteriol. 85, 1274–1279 (1963).

32. Kirkcaldy, R. D. Neisseria gonorrhoeae Antimicrobial Susceptibility Surveillance — The Gonococcal Isolate Surveillance Project, 27 Sites, United States, 2014. MMWR Surveill. Summ. 65, (2016).

33. Mukherjee, A. et al. Fluoroquinolone resistance-conferring gyrA variants alter the fitness cost and potentiate the resistance of the zoliflodacin resistance mutation gyrBD429N in Neisseria gonorrhoeae. 2026.05.04.722797 Preprint at 10.64898/2026.05.04.722797 (2026).

34. Odds, F. C. Synergy, antagonism, and what the chequerboard puts between them. J. Antimicrob. Chemother. 52, 1 (2003).

35. Singh, V., Bala, M., Bhargava, A., Kakran, M. & Bhatnagar, R. In vitro efficacy of 21 dual antimicrobial combinations comprising novel and currently recommended combinations for treatment of drug resistant gonorrhoea in future era. PLoS ONE 13, e0193678 (2018).

36. Torella, J. P., Chait, R. & Kishony, R. Optimal Drug Synergy in Antimicrobial Treatments. PLOS Comput. Biol. 6, e1000796 (2010).

37. Hegreness, M., Shoresh, N., Damian, D., Hartl, D. & Kishony, R. Accelerated evolution of resistance in multidrug environments. Proc. Natl. Acad. Sci. 105, 13977–13981 (2008).

38. Foerster, S. et al. In vitro antimicrobial combination testing of and evolution of resistance to the first-in-class spiropyrimidinetrione zoliflodacin combined with six therapeutically relevant antimicrobials for Neisseria gonorrhoeae. J. Antimicrob. Chemother. 74, 3521–3529 (2019).

39. Jacobsson, S. et al. Pharmacodynamics of zoliflodacin plus doxycycline combination therapy against Neisseria gonorrhoeae in a gonococcal hollow-fiber infection model. Front. Pharmacol. 14, 1291885 (2023).

40. Jensen, J. S., Nørgaard, C., Scangarella-Oman, N. & Unemo, M. In vitro activity of the first-in-class triazaacenaphthylene gepotidacin alone and in combination with doxycycline against drug-resistant and -susceptible Mycoplasma genitalium. Emerg. Microbes Infect. 9, 1388–1392 (2020).

41. Yechezkel, M. et al. Rapid loss of doxycycline effectiveness against gonorrhea after implementation of post-exposure prophylaxis in southern California: an observational study. 2026.01.07.26343623 Preprint at 10.64898/2026.01.07.26343623 (2026).

42. Do, K., Unemo, M., Kenyon, C., Hocking, J. S. & Kong, F. Y. S. Tetracycline-resistant Neisseria gonorrhoeae global estimates—impacts on doxycycline post-exposure prophylaxis implementation and monitoring: a systematic review. JAC-Antimicrob. Resist. 7, dlaf120 (2025).

43. Jacobsson, S. et al. High prevalence of tetracycline resistance in Neisseria gonorrhoeae across 22 European countries, 2024. Eurosurveillance 31, 2500965 (2026).

44. Igo, M. et al. A metagenomic analysis for combination therapy of multiple classes of antibiotics on the prevention of the spread of antibiotic-resistant genes. Gut Microbes 15, 2271150 (2023).

45. Bristow, C. C. et al. Whole-Genome Sequencing to Predict Antimicrobial Susceptibility Profiles in Neisseria gonorrhoeae. J. Infect. Dis. 227, 917–925 (2023).

46. Eyre, D. W. et al. Detection in the United Kingdom of the Neisseria gonorrhoeae FC428 clone, with ceftriaxone resistance and intermediate resistance to azithromycin, October to December 2018. Euro Surveill. 24, 1900147 (2019).

47. Swanson, J. Studies on gonococcus infection. II. Freeze-fracture, freeze-etch studies on gonococci. J. Exp. Med. 136, 1258–1271 (1972).

48. Peng, J.-P. et al. A Whole-genome Sequencing Analysis of Neisseria gonorrhoeae Isolates in China: An Observational Study. eClinicalMedicine 7, 47–54 (2019).

49. Mortimer, T. D. et al. The Distribution and Spread of Susceptible and Resistant Neisseria gonorrhoeae Across Demographic Groups in a Major Metropolitan Center. Clin. Infect. Dis. 73, e3146–e3155 (2021).

